# Phages prevent biofilm formation on catheters under flow

**DOI:** 10.1101/2023.07.26.550655

**Authors:** Hoda Bseikri, Slawomir Michniewski, Eduardo Goicoechea Serrano, Eleanor Jameson

## Abstract

Biofilms pose a significant challenge in medical settings, leading to persistent infections. Phage therapy has shown promise in biofilm eradication, but its effectiveness under dynamic flow conditions remains unclear. Here we use two novel phages isolated on *Klebsiella*, Llofrudd and Samara, and characterized their genomes, host range and virulence. In this study, we built a simple catheterised bladder model with flow to investigate the impact of phage treatment on biofilm viability in a flow-based catheter model. Our analyses demonstrate that phages Llofrudd and Samara are the same species and infect a limited number of strains (3/222), but across three species: *Klebsiella aerogenes, Klebsiella pneumoniae* and *E. coli*. Phage treatment significantly reduced *E. coli* biofilm viability in catheters both in static conditions and under flow, highlighting the potential of phage therapy as an intervention strategy for catheter associated urinary tract infections (CAUTI).

## Introduction

The use of indwelling catheters is common in health care settings, yet there is a risk of catheter associated urinary tract infections (CAUTIs) (*Nicolle, 2014*). Approximately 86% of hospital patients undergoing operations require indwelling catheters, with the cost of catheter use, management and CAUTIs estimated to be £1-2.5 billion per annum in the National Health Service (*Feneley et al., 2015*). CAUTIs accounted for >17% of healthcare acquired infections making them the third most common infection in these settings (*Zarb et al., 2012*). CAUTIs can develop into life-threatening complications such as pneumonia, bacteraemia or sepsis (*Jacobsen et al., 2008*), and are associated with extended stays (*Green et al., 1982*) and medical costs (*Hollenbeak and Schilling, 2018*). Around 1-4% of CAUTIs develop into bacteraemia, which has a fatality rate of ∼13 % (*Bryan and Reynolds, 1984*). The use of indwelling catheters is common practice in the medical field, with long-term catheters being used for up to three months (*Hall et al., 2020*). However, prolonged catheter use risks bacteria forming biofilms on the catheter, which increases the risk of patients developing a CAUTI (*Al-Hazmi, 2015; Clayton, 2017; Cortese et al., 2018*).

A further complication of bacterial infections is antimicrobial resistance (AMR), which is increasing and poses a significant global threat to public health (*O’neill, 2014; Zarb et al., 2012*). In 2019, ∼ 4.8 million deaths were associated with AMR (*Murray et al., 2022*), and hospital-associated AMR infections and deaths rose a further 15% during 2020 (*Centers for Disease Control and Prevention, 2022*). The two most prevalent causes of CAUTIs, *Escherichia coli (E. coli)* and *Klebsiella* (*Gul et al., 2021*), have been identified as key priority pathogens for the development of new antimicrobials by the World Health Organization due to widespread AMR associated with them (*Tacconelli et al., 2018, 2017*). Both *E. coli* and *Klebsiella* readily form biofilms on catheters, making them even more recalcitrant to antimicrobials (*Hidron et al., 2008; Jacobsen et al., 2008; Sabir et al., 2017; Wazait et al., 2003*). These biofilms can form under different nutrient conditions and flow levels, and once formed are hard to remove (*Melo and Vieira, 1999*). The increasing prevalence of AMR and biofilm formation in CAUTIs makes the need for alternative or complementary treatments to existing antibiotics critical (*Gul et al., 2021; Marantidis and Sussman, 2023*). Bacteriophages (phages for short) represent an alternative approach with the potential to treat or prevent AMR bacterial infections, including CAUTIs (*Chegini et al., 2021; Lehman and Donlan, 2015; Milo et al., 2017; Townsend et al., 2020*).

Phage therapy has been successfully used to treat patients with a range of bacterial infections with little or no adverse side effects (*Bruttin and Brussow, 2005; El Haddad et al., 2019*). This illustrates the promise of phages for medical applications, but they are not yet a standard treatment and more focussed research is required for a variety of infections, including urinary tract infections (UTIs). Studies have demonstrated that phages can be readily isolated against UTI causing bacteria and have potential to treat and prevent UTIs (*Chegini et al., 2021; Lehman and Donlan, 2015; Melo et al., 2016; Milo et al., 2017; Townsend et al., 2020*). One mode of delivery is to coat catheters with phages, as a prophylaxis against the most common and problematic CAUTIs (*Curtin and Donlan, 2006; Milo et al., 2017; Townsend et al., 2020*). For phages to be successfully used in catheters we need to understand how well they perform in the conditions experienced in catheters; previously phages have been tested on catheter material and in artificial urine (*Townsend et al., 2020*). The efficacy of phages in dynamic flow conditions, such as shear-stress of urine flow encountered in catheters, remains poorly understood (*MacCallum et al., 2015; Ramstedt et al., 2019*). In this study we developed and tested a simple, cheap bladder model with the aim to investigate if phages can reduce CAUTI biofilm viability in a flow-based catheter model.

## Materials and Methods

### Phage strains

The phages used in the study were the closely related phages vB_KaS-Llofrudd and vB_KaS-Samara (the phages will be referred to as Llofrudd and Samara respectively from this point forwards). Both phages were isolated from cow slurry on *Klebsiella aerogenes* DSM 30053. The phages have a siphovirus-like morphology. Llofrudd and Samara were chosen for their efficacy against both the *Klebsiella* host of isolation and a tractable non-pathogenic reference strain of *E. coli* as a test case to establish the model system.

### Genome sequencing and annotation

Phage genomes were sequenced by MicrobesNG (Birmingham, UK) using Illumina sequencing platform. Trimmed reads were assembled de-novo using SPAdes v3.10.1 (*Bankevich et al., 2012*). Assembled phage genomes were annotated with Prokka v1.14.6 (*Seemann, 2014*) using the PHROGS database (*Terzian et al., 2021*). The genomes of Llofrudd and Samara were compared against INPHARED (*Cook et al., 2021*) database of known phage genomes to identify their closest relatives using MASH (*Ondov et al., 2016*) with “dist -d 0.15” option. Subsequently, their intergenomic similarities were calculated and putative species and genera clusters were determined using VIRIDIC (*Moraru et al., 2020*).

### Bacterial strains and host range

Spot testing was carried out against 148 *Klebsiella* spp strains, 125 of which were of clinical origin and 23 reference strains from commercial collections (the species included were *K. aerogenes, K. pnuemoniae, K. oxytoca, K. variicola, K. michiganensis and K. quasipneumoniae*), and 74 *E. coli*, 71 of which were of clinical origin and 3 reference strains, to determine the host range of phages Llofrudd and Samara, following the method described previously (*Townsend et al., 2021*). Briefly phage stock solutions were serial diluted to 1 × 10^−7^, and 5 µL of each dilution was spotted onto a 0.4% overlay agar bacterial lawn, followed by overnight incubation at 37 °C. Zones of bacterial lawn clearing, indicating cell lysis, were recorded with particular attention to visible plaques.

The bacteria carried forwards for use in the biofilm experiments is the non-pathogenic bacteria *Escherichia coli* K12 MG1655 (referred to as *E. coli* K12 from this point on)(*Blattner et al., 1997*).

### Phage virulence index

The virulence index (VP) of phages Llofrudd and Samara was calculated (*Storms et al., 2020*). A high titre phage lysate was prepared and used as reference throughout the study. Briefly, *E. coli* K12 cultures in the exponential growth phase were adjusted by dilution to an optical density equivalent to 1 × 10^8^ colony forming units/ml (CFU/mL) in LB. Phages were serially diluted in a 96 well plate from 1 × 10^8^ to 10 plaque forming units/ml (PFU/ml) in 100 µL volumes. To this 100 µL of the adjusted *E. coli* K12 culture was added, resulting in multiplicity of infections (MOIs) from 1 to 10^−7^. MOI is a parameter that describes the phage to bacteria ratio. The 96 well plate was transferred to the plate reader at 37 °C with shaking, the optical density of each well was read at 600 nm every 5 min for 18 h.

The area under the curve of the resulting growth curves at each MOI was calculated from infection to the exponential growth stage. For each MOI the VP was calculated following the method (*Storms et al., 2020*) in RStudio (version 1.1.463). VP is a measure of the phage virulence against a bacterium, on a 0–1 scale (0 = no impact on bacterial growth, to 1, instant complete killing).

### Phage quantification

Phages were quantified by plaque assays as previously described (*Townsend et al., 2021*). Briefly, serial dilutions of the phages were made. Then 50 μl of each phage dilution was mixed with 500 μl of logarithmic growth phase *E. coli* K12 (approximately OD600nm 0.2) and incubated at room temperature for 10 min. To this mix 2.5 mL of hand-warm, molten LB agar (0.5% weight/volume, supplemented with 5 mM CaCl_2_) was added and mixed by swirling. Mixed agar was overlayed onto 1% LB agar plates. After setting the overlay agar plates were inverted and incubated at 37°C overnight. Plaques were counted and the titre of was presented as the number of PFU/ml.

To quantify plaques from catheters the plaque assay was adapted. For this, 1 cm long catheter segments were incubated with 500 μl logarithmic phage *E. coli* K12 for 10 min. The catheter segments were removed, 2.5 mL of hand-warm, molten LB agar (0.5%) was added and mixed by swirling. The above procedure for making the overlay plates was then followed.

### Biofilm growth in static 96 well plates

Biofilms grown in cell culture plates were used as a simple, *in vitro* model to assess the ability of *E. coli* K12 to form biofilms and the ability of phages to inhibit biofilm formation and biofilms. *E. coli* cultures were standardised to 1 × 10^6^ CFU/mL in LB. Biofilms were grown in 96-well flat bottom cell culture treated plate (CytoOne Plate, Starlab, UK), 100 μL of standardised *E. coli* culture was added. For the treatments 100 μl of phage lysate at MOIs of 0.1, 1 and 10 was added to the *E. coli* culture. To produce the phage stocks at different MOIs, high titre phage lysates were diluted in LB media (i.e. the phage stock for MOI 0.1 = 1 × 10^5^ PFU/mL, MOI 1 = 1 × 10^6^ PFU/mL and MOI 10 = 1 × 10^7^ PFU/mL). For the treatments, biofilms were grown in the presence of either phage Llofrudd and Samara, positive controls with no phage treatment and negative controls containing phage alone, were included. To form a biofilm, the plates were incubated statically for 24 h at 37 °C. After 24 h, biofilm viability had stabilised and considered mature. All tests were carried out in technical triplicate, on three occasions for biological replicates.

### Biofilm growth in static catheters

The catheters used in this study were Foley silicone catheters, 18fg 5/30ml balloon urinary catheters (Medisave). To grow biofilms in catheters under static conditions two methods were used.

The protocol for the growth of biofilm on catheters under static conditions was based on the method we previously described (*Townsend et al., 2020*). Briefly, 1 cm catheters segments were incubated aerobically in 10 ml LB cultures *E. coli* at OD 0.6 for 2 hr with shaking. These catheters were then transferred to 24 well culture plates containing 2ml of LB media and incubated overnight at 37 °C. For the phage treated group, 100 μl of phage lysates 1 × 10^6^ pfu/ml was added to the wells. The negative control contained phage lysates alone. All tests were carried out in technical triplicate, with three biological replicates.

### Catheterised bladder model

A simple catheterised bladder model was constructed using readily available lab consumables, as shown in Figure 1. To represent the bladder a 50 ml falcon tube was used. This bladder is fed with media through a media inlet consisting of a silicone tube inserted into the lid of the flacon tube. A 10 cm foley catheter segment (18fg 5/30ml balloon urinary catheters, Medisave) was inserted through a hole in the bottom of the falcon tube, and protruded 5 cm into the falcon tube. At the 35 ml graduation on the falcon tube a side inlet was inserted, consisting of a 5 cm segment of silicone tube to allow the addition of bacteria and phages to the catheterised bladder, this was clamped when not in use. All inlets and outlets were drilled, and tubing or catheter segments were inserted before autoclave sterilisation. Following sterilisation the tubing was glued in place using a hot melt adhesive (Power Adhesives 240-12-300-CRP-TP16-RS).

**Figure 1.**
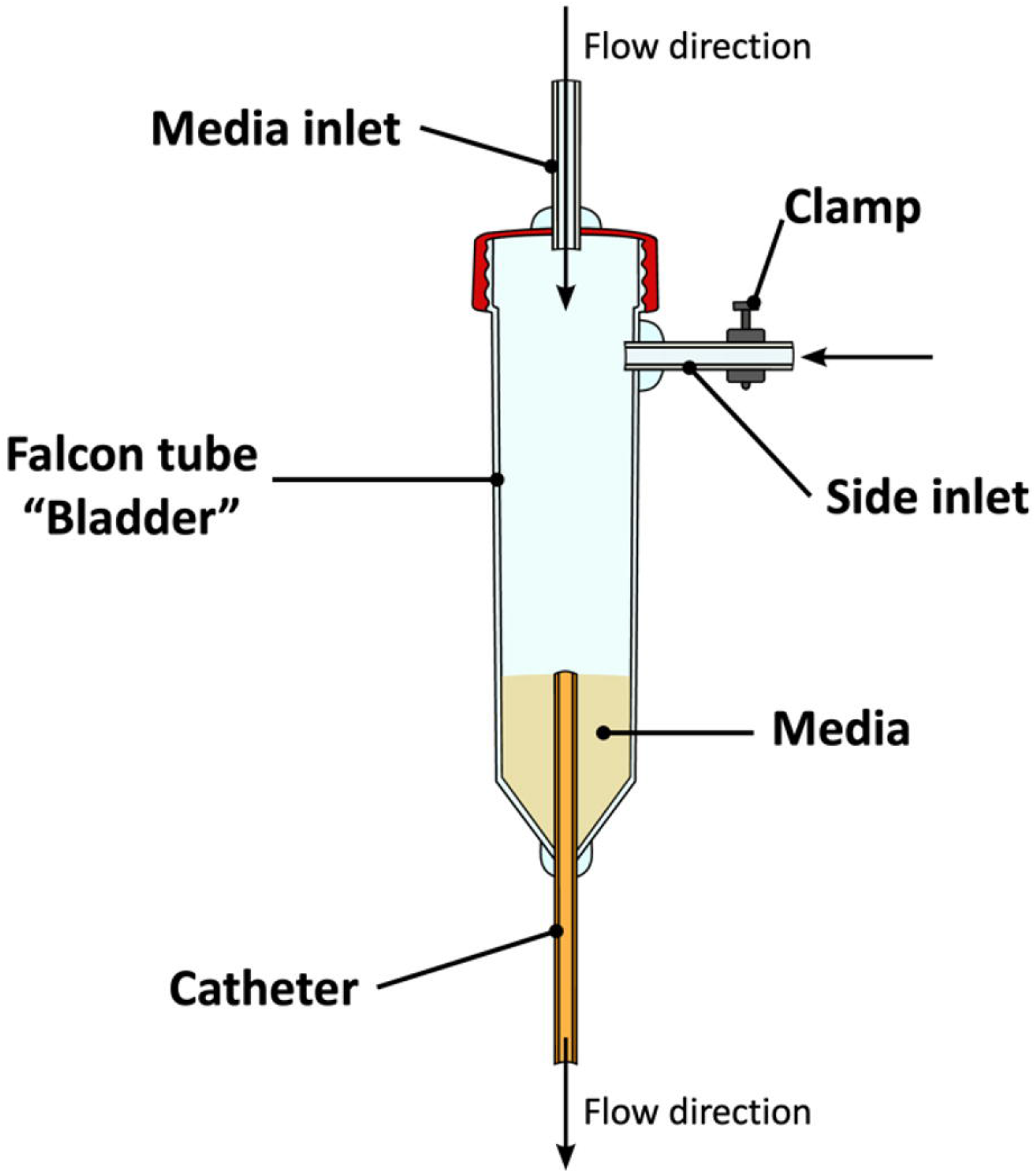
Diagram of the catheterised bladder model, used to test phage treatments on bacterial biofilms in urinary catheters under flow. Media was added by continuous flow at 3ml/min (as indicated by the flow direction arrow) through the media inlet, the side inlet was used for the addition of bacteria and phage, the catheter at the base of the falcon tube allows drainage of the media through the catheter to waste.

The catheterised bladder model shown in figure 1 allows the falcon tube bladder to fill with media until it covers the top of the catheter before media will flow through the catheter, mimicking the emptying of a bladder. The bottom of the falcon tube with the catheter segment was placed in a 1 L waste bottle and sealed with parafilm to maintain sterile conditions.

### Under-flow biofilms

Under-flow biofilms were grown in catheter segments in the catheterised bladder model (fig. 1). Prior to inoculation LB media was pumped through the system for 1 hr at a rate of 3 ml/min. Bacterial biofilms were established as follows: the catheter was filled with 20 ml *E. coli* K12 inoculum in LB at 1 × 10^6^ CFU/ml. The inoculum was incubated in the catheter for 2 hr under static conditions to establish a colonising biofilm. For the treatments, following static bacterial incubation, 20 ml of filtered phage lysate at 1 × 10^6^ PFU/ml or LB media (for the no phage control) was injected through the side inlet of the bladder model. The phages were incubated for 10 min, under static conditions to enable phage-bacteria interactions (mimicking a phage treatment or coating on the catheters). Initiating LB media flow through the bladder model and catheter. Flow rate was maintained at 3 ml/min using a 323S Drive peristaltic pump (Watson-Marlow Ltd). Phages Llofrudd and Samara were applied as treatments in parallel catheterised bladder models. All under-flow biofilms were grown for 24 hrs. All tests were carried out in technical quadruplet, with three biological replicates.

### Biofilm viability analysis

To quantify actively respiring cells in biofilms, the resazurin assay was used. Before carrying out the assay media was removed from the biofilms by pipetting, then the biofilm was washed twice with phosphate buffered saline. An aliquot (200 µl in 96 well plates or 2 ml for catheter sections) of 10 mg/ml resazurin solution was added to the biofilms and then incubated at 37 °C. The end point was determined when the positive control (*E. coli* K12 only) changed from blue to pink. When the end point was reached the absorbance of the resazurin solution was measured at 570 nm and 600 nm to calculate the percentage biofilm viability.

For the under-flow catheter biofilm experiments; following 24 hr of biofilm growth in catheterised bladder model, the catheter was extracted and cut into 1 cm sections to assess biofilm viability, both 1 cm end sections were discarded due to variability in biofilm formation.

## Results

### Host range

The host range was generated against the clinical and reference panel. Both phages Llofrudd and Samara infected and produced plaques in 2/148 *Klebsiella* spp. and 1/74 E. coli. The strains were: reference strain *Klebsiella aerogenes* 30053, clinical isolate *K. pneumoniae* WALES42 and reference strain *E. coli* K12. The reference strain *E. coli* K12 was selected as the host for all experiments from this point forwards because it is non-pathogenic, which was important when developing a flow-based system due to potential containment and spill risks.

### Phage virulence index

The virulence index for Llofrudd was 0.54, and Samara had a value of 0.56.

### Sequencing

The reconstructed genome size of Llofrudd was 49,702 bp and Samara was 49,646 bp. Genomes have been uploaded to the deposited in the ENA under the accessions OY639338 and OY639337, respectively. Both phages Llofrudd and Samara showed >99.9 % MASH similarity to each other and to vB_KaS-Ahsoka (accession LR881108), vB_KaS-Gatomon (LR881110) and vB_KppS-Samwise (LR881107), and >79 % MASH similarity to Escherichia phage Henu7 (MN019128), Enterobacter phage N5822 (MW032452), Escherichia phage IsaakIselin (MZ501077) and Escherichia phage JohannLBurckhardt (MZ501085). VIRIDIC analysis results at 70 % intergenomic similarity cutoff indicated that these phages belong to the same genus: *Henuseptimavirus* (Family *Drexlerviridae*, Subfamily *Tempevirinae*). However, only phages Llofrudd, Samara, Gatomon, Ahsoka and Samwise were clustered together at >95% intergenomic similarity, indicating that they belong to a separate, yet unclassified phage species in the defined *Henuseptimavirus* genus represented by Henu7 (table1) (*Moraru et al., 2020*).

**Table 1.**
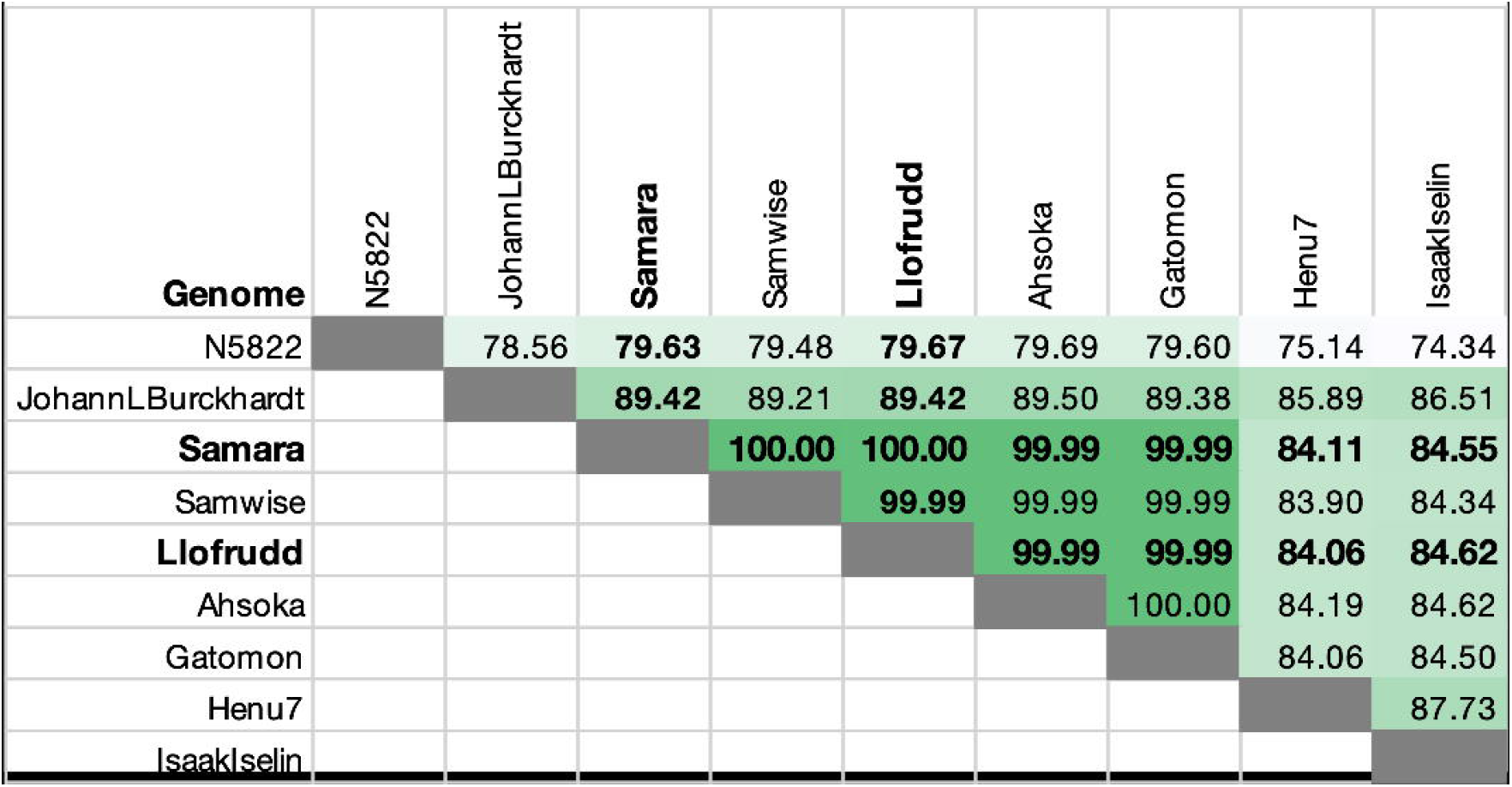
Phage ANI similarity to the closest sequenced phage genome available calculated by VIRIDIC (*Moraru et al., 2020*).

### Phage treatment of biofilms grown under static conditions

Phages reduced biofilm viability in both 96 well plates and on static catheters (Fig. 2). In the 96 well plate grown biofilms, the biofilms grown with both phages Llofrudd and Samara had lower viability than the no phage controls. When phages were supplied at an MOI = 10, the highest concentration of phage to bacteria a small decrease from 81 % in the control to 64 % with Llofrudd and 50 % with Samara was observed (Fig 2A). At lower MOI of 1 and 0.1 a greater decrease with phage treatments were observed, ∼ 30 % biofilm viability was observed for all the phage treatments, compared to the control ∼ 80 % (Fig S1A). A similar pattern was observed on the statically incubated catheter segments, biofilm viability was lower in the phage treatments < 40 % for the phage treatments, compared to the control ∼ 80 % (Fig S1B).

**Figure 2.**
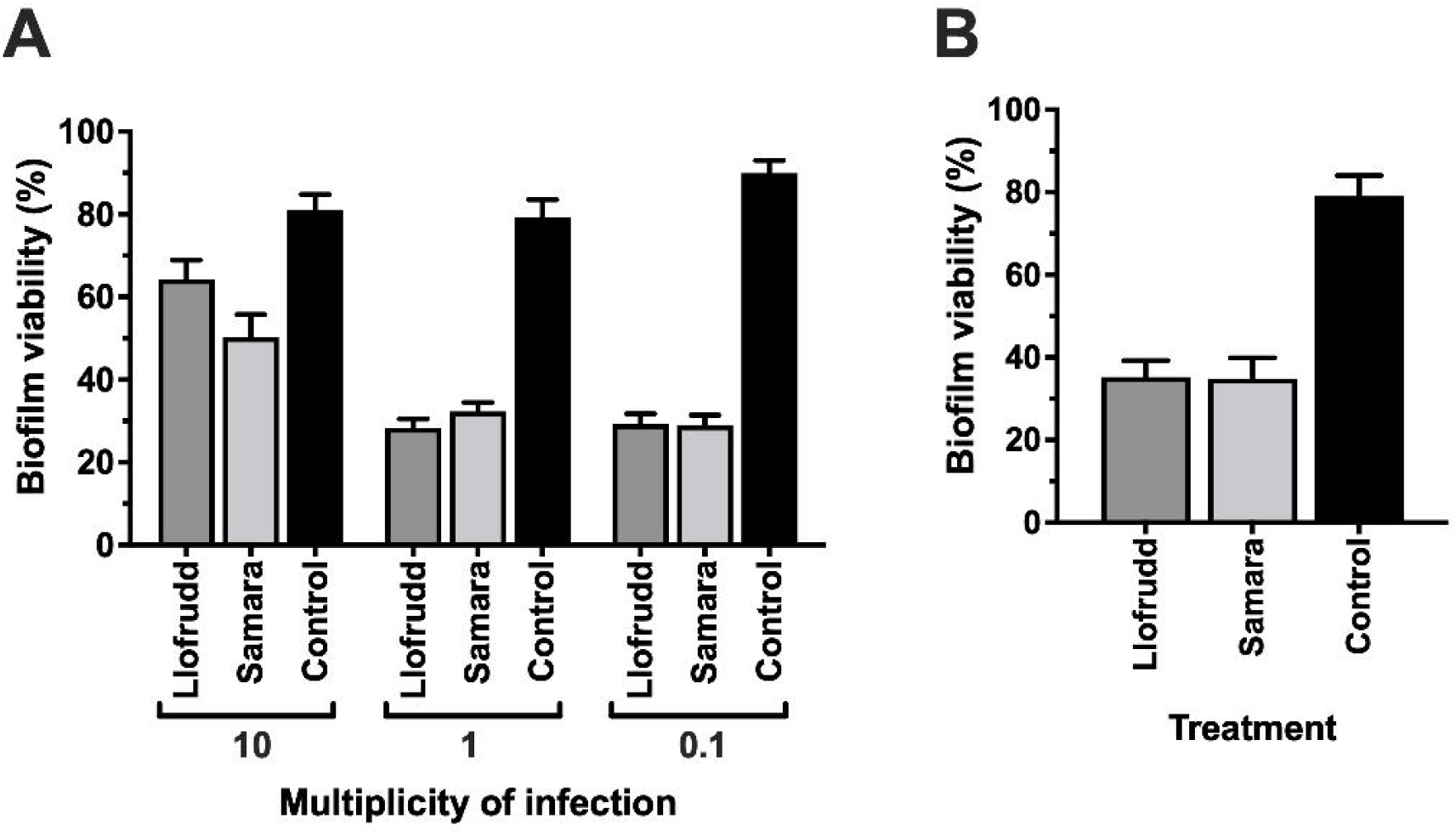
Phage treatments reduce the biofilm viability of *E. coli* K12 under static conditions: **A.** Grown in 96 well plates; the phages vB_KaS-Llofrudd (Llofrudd) and vB_KaS-Samara (Samara) were added at the beginning of biofilm formation at multiplicity of infections (MOI) 10, 1 and 0.1, as indicated on the X-axis. **B.** Grown on foley catheters; phages Llofrudd and Samara were added at the beginning of biofilm formation at a concentration of 1 × 10^6^ PFU/ml. The first two grey bars represent phage addition treatments, either phage Llofrudd or Samara the third black bar represents *E. coli* K12 with no phage added (No phage control). All biofilms were grown for 24 hr at 37 °C under with or without phage treatments. Bars represent the mean of three biological replicates, in technical triplicate and error bars represent the standard error of mean.

### Flow model

The flow model was successfully run for three independent experiments that formed the biological triplicates. Flow was maintained throughout the experiment and a biofilm was able to form. The variation in biofilm viability observed in the catheter segments was higher than observed in the static catheter experiments, indicating that the shear force created by the continuous flow of media may have impacted biofilm distribution (fig. 3).

**Figure 3.**
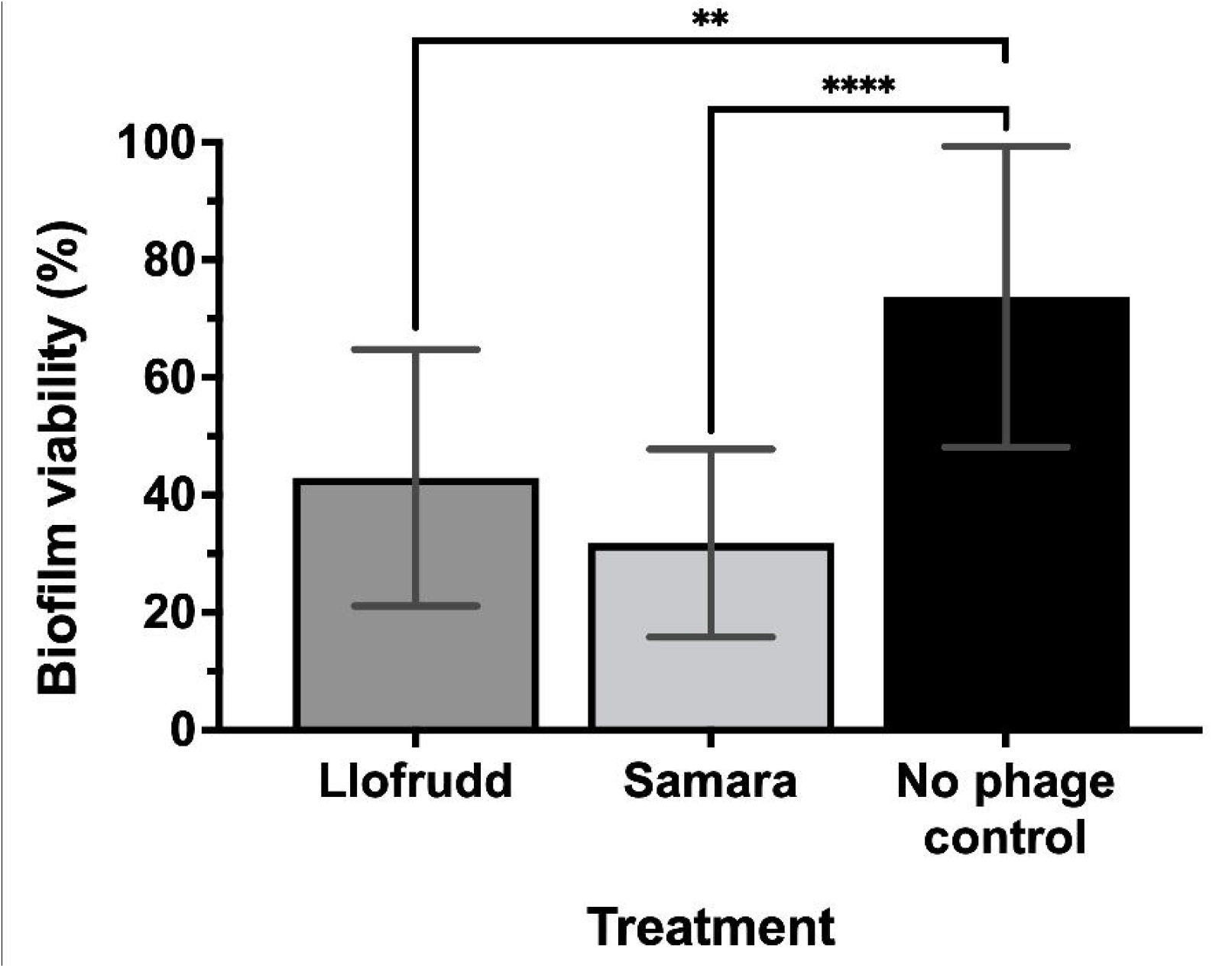
Phage treatments reduce the biofilm viability of *E. coli* K12 grown under flow in the catheterised bladder model. The biofilms were grown for 24 hr at 37 °C under a continuous flow rate of 3 ml/min with or without phage treatments. The first two grey bars represent phage addition treatments, using either vB_KaS-Llofrudd (Llofrudd) or vB_KaS-Samara (Samara) as labelled, the third black bar represents *E. coli* K12 with no phage added (No phage control). Brackets with the associated symbols denote significant differences (t-test) compared to the positive control, ** denotes p value ≤ 0.01, **** p value ≤ 0.0001. Bars represent the mean of three biological replicates, with four technical replicates and error bars represent the standard error of mean.

### Phage treatment of the catheterised bladder model biofilms under-flow

Phages significantly reduced biofilm viability on catheters where flow was introduced (fig. 3). The catheters with the addition of either phage Llofrudd or Samara had statistically lower viability than the no phage controls; ∼ 40 % biofilm viability was observed when phage Llofrudd was added, this figure was even lower for Samara ∼ 30 % viability, compared to the no phage controls that had ∼ 70 % viability (fig 3).

## Discussion

We observed that phage treatment with Llofrudd and Samara significantly reduced biofilm viability, with a greater reduction seen in static conditions (50 % reduction) compared to under continuous flow with a bladder reservoir (30 % reduction). This supports a previous study that have demonstrated the ability of phages to prevent biofilm formation under flow (*Lehman and Donlan, 2015*). However, our findings show that flow affected the extent of biofilm-reduction by phages, indicating the importance of considering the role of urine flow dynamics when optimising phages application for preventing and treating catheter-associated urinary tract infections (CAUTIs). Urine flow can influence the distribution of bacteria and biofilms along the catheter surface (*Ganderton et al., 1992; Melo and Vieira, 1999*). Shear forces generated by flow can disrupt biofilm formation (*Tsagkari et al., 2022*) and potentially enhance the accessibility of phages to target bacterial cells within the biofilm matrix. Our study observed more variation in biofilm viability in catheters under flow than static conditions, indicating the potential impact of flow on biofilm distribution. These findings suggest that phage therapy, combined with appropriate urine flow management, could be an effective strategy to prevent and treat CAUTIs, reducing patient discomfort, hospitalization duration, and healthcare costs.

Furthermore, our study highlights the ability of phages isolated on *Klebsiella aerogenes* to infect and prevent biofilm formation in another genera, *Escherichia*. Although we have only used single phages in this study, it indicates that the application of *Klebsiella* phages to prevent CAUTI, may also reduce *E. coli* infections. Our previous work has demonstrated the importance of using phage cocktails rather than single phage treatments (*Townsend et al., 2020*). The combination of multiple phages, with different specificities could enhance the overall efficacy of phages under flow in catheters (*Lehman and Donlan, 2015*). It will be important to run tests for longer (1 - 7 days, the cut off for short-term catheter use (*Al-Hazmi, 2015; Hall et al., 2020*) to optimise dose and application methods for phages, particularly as low concentrations of phage treatment can initiate biofilm proliferation through stress induced systems in host bacteria (*Fernandez et al., 2017*).

A simple bladder model built from cheap, readily available laboratory supplies is and effective way to test CAUTIs *in vitro* and introduce flow. Our study demonstrates its applicability for testing antimicrobials in catheters to prevent CAUTIs. The model can be housed in existing incubators and needs very little equipment to study the impact of flow dynamics on catheter biofilms in the context of treatment efficacy. By utilizing our bladder model and building upon these findings, researchers can develop more effective strategies for preventing and managing UTIs, ultimately improving patient outcomes. This simple bladder model allowed us to investigate the interplay between flow, phage treatment, and biofilm viability, to provide insights into the development of effective clinical interventions.

## Conclusions

Our study demonstrates the effectiveness of phage treatment in reducing biofilm viability in both static and flow-based catheter models. Our bladder model provides a practical tool for researching CAUTI, allowing investigation of factors influencing biofilm development and treatment. Phages Llofrudd and Samara, despite the small number of hosts they infect can infect different species and genera. This underscores the importance of considering flow dynamics and phage choice on CAUTI biofilm formation. Our study highlights the value of phage therapy and the bladder model in advancing CAUTI research and driving innovation in UTI management.

## CRediT authorship contribution statement

**Hoda Bseikri:** Methodology, Investigation, Writing – original draft, Visualization. **Slawomir Michniewski:** Sequencing, Bioinformatics, Writing - review & editing. **Eduardo Goicoechea Serrano:** Phage characterisation, Funding acquisition. **Eleanor Jameson:** Conceptualization, Writing, Visualization, Supervision, Funding acquisition.

## Acknowledgements

This work was supported by a Warwick Integrative Synthetic Biology (WISB) early career fellowship, funded jointly by BBSRC/EPSRC, grant ref: BB/M017982/1 under the UK Research Councils’ Synthetic Biology for Growth programme, and the Warwick Flexible Talent Mobility Award, funded by BBSRC grant ref: BB/S507982/1.

